# A pipeline for assessing the quality of images and metadata from crowd-sourced databases

**DOI:** 10.1101/2022.04.29.490112

**Authors:** Jackie Billotte

## Abstract

Crowd-sourced biodiversity databases provide easy access to data and images for ecological education and research. One concern with using publicly sourced databases; however, is the quality of their images, taxonomic descriptions, and geographical metadata. The method presented in this paper attempts to address this concern using a suite of pipelines to evaluate taxonomic consistency, how well geo-tagging fits known distributions, and the image quality of crowd-sourced data acquired from iNaturalist, a crowd-sourced biodiversity database. Additionally, it provides researchers that use these datasets to report a quantifiable assessment of the taxonomic consistency. The pipeline allows users to analyze multiple images from iNaturalist and their associated metadata; to determine the level of taxonomic identification (family, genera, species) for each occurrence; whether the taxonomy label for an image matches accepted nesting of families, genera, and species; and whether geo-tags match the distribution of the taxon described using occurrence data from the Global Biodiversity Infrastructure Facility (GBIF) as a reference. Additionally, image quality is assessed using BRISQUE, an algorithm that allows for image quality evaluation without a reference photo. Entries from the order Araneae (spiders) are used as a case study. Overall, the results suggest that iNaturalist can provide large metadata and image sets for research. Given the inevitability of some low-quality observations, this pipeline provides a valuable resource for researchers and educators to evaluate the quality of iNaturalist and other crowd-sourced data.

## Introduction

In the past ten years, biodiversity, conservation, and taxonomical research have increasingly utilized community science initiatives, including virtual platforms of user-uploaded biodiversity observations (Cull, 2021; Mesaglio & Callaghan, 2021; Ryan et al., 2018). These large, crowd-sourced biodiversity databases contribute to scientific advancement, with posted observations providing evidence of unknown species, surveillance of field sites, and newly documented animal behaviors (Wilson et al., 2020). For animal groups that are difficult to locate in the wild or whose collection would prove too time-consuming and costly, crowd-sourced observations could provide information on species ranges or invasions. For example, the first occurrence of Pseudeuophrymings (Araneae: Salticidae: Euophryini), outside its native range was documented by non-experts on biodiversity databases (Kaldari, 2019). Since observations from crowd-sourced data are usually accompanied by geographic information, erratic comparison of these observations against known ranges can help highlight observations made well outside the known range. Streamlining the use of free and crowd-sourced biodiversity databases could make the detection of new species faster and more efficient (Michonneau & Paulay, 2014; “What Is GBIF?” 2021). iNaturalist, one such crowd-sourced biodiversity database, provides a large platform for gathering global biodiversity data. Founded in 2008 by Nate Agrin, Jessica Kline, and Ken-ichi Ueda as part of a graduate school project for the University of California, Berkeley School of Informatics Master’s project (Agrin et al., 2014), iNaturalist has grown in popularity, and now contains more than 68 million observations. Using iNaturalist and its accompanying mobile app, users can post images of organisms and provide a geo-tagged location for that observation. Once a user posts an image, an image-based machine learning algorithm suggests a taxonomic identification. Users can then vote in agreement with that identification or suggest an alternative identification until one proposed taxonomy receives enough votes for general acceptance. The program labels observations with agreed-upon taxonomic identification and higher quality images as “Research grade” (Agrin et al., 2014). Like iNaturalist, the Global Biodiversity Information Facility (GBIF) is an open-access infrastructure that provides observational data of species using a range of published sources (Michonneau & Paulay, 2014; “What Is GBIF?,” 2021). The observation records from GBIF come from participating institutions and publications. The curated data are available publicly, but unlike iNaturalist, contributions are limited to only approved organizations, making the data, in theory, more robust, but also more limited (“What Is GBIF?” 2021).

Image and metadata quality remains a concern when using crowd-sourced data (Cull, 2021; Moudrý & Devillers, 2020). Though sites implement some quality control, the volume of entries prevents more detailed scrutiny of the data. If the data are of high enough quality, iNaturalist could be used as a digital alternative to physical collections for rigorous morphological comparisons (Mugford et al., 2021). These tasks would require consistently labeled images, no duplicate entries, correct geo-tags, and image quality high enough to permit accurate identification and comparison (Hochmairid et al., 2020). Currently, researchers and educators must evaluate entry quality manually. Given the millions of observations recorded for any given taxonomic group, this process is time and resource intensive and represents the biggest barrier to using crowd-sourced data. This study seeks to address this barrier by providing a pipeline to quantify and assess the quality of data currently available on iNaturalist using the order Araneae as a case study (Figure 1). Much like a physical pipeline, pipelines allow for the output of the computing process to automatically feed into the next process, allowing information to flow in one direction.

**Figure 1.**
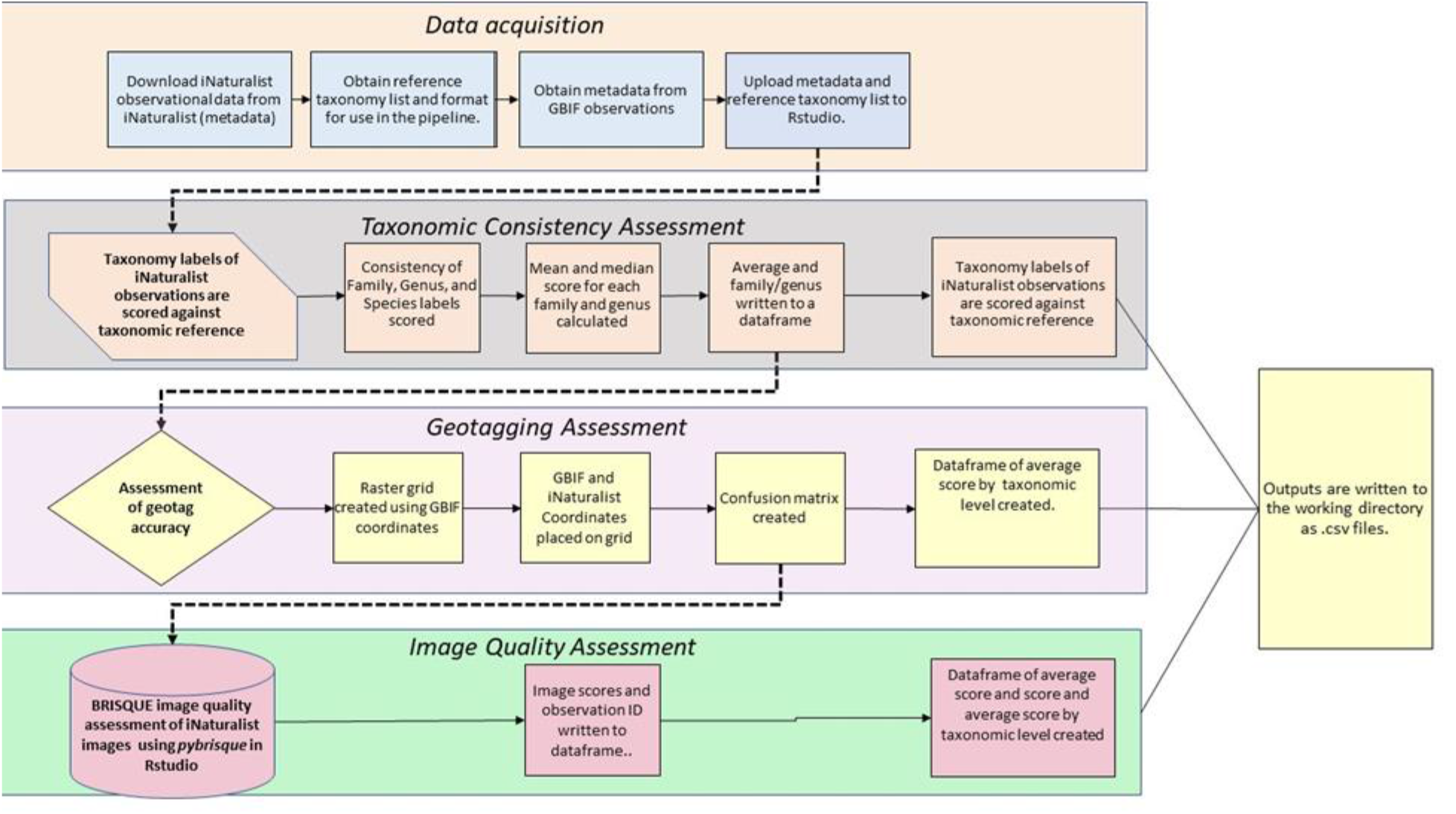
Workflow for pipeline suite. Data is acquired from iNaturalist and GBIF. The first pipeline compares the taxonomic consistency of the iNaturalist data. The second assesses the quality of the images using pyBRISQUE, and the third compares the geo-tagging from the iNaturalist observations to those obtained by GBIF. Bold dashed lines represent transitions between pipelines in the suite of pipelines. Data frames and assigned variables will be passed between pipelines. All outputs are saved to the working directory as .csv files.

Fully assessing datasets acquired from iNaturalist (and similar databases) can illuminate areas that may still need to improve and how the research community may create a more accurate and robust database.

## Material and methods

### Data acquisition

I applied the pipeline to observations from iNaturalist for the order Araneae as a case study. Spiders constitute a valuable, yet understudied order of invertebrates (Nyffeler et al., 1994; Schwerdt et al., 2018). Difficulties associated with locating, observing, and collecting spider specimens partly account for the deficit of data on species ranges and diversity, and the lack of information on habitat and behavior (Cardoso et al., 2011). Community science initiatives (projects that partner researchers with community volunteers) help researchers expand arachnid research while also providing public outreach and education (Cull, 2021). While this pipeline was originally meant to be used with iNaturalist, its potential use with other databases necessitates acquiring images and their accompanying metadata before using the pipeline, as each database varies on how users access data. For the Araneae case study, I searched for and downloaded observations for each family under the order Araneae on iNaturalist on July 21, 2021 (“iNaturalist,” 2014). I then searched for and downloaded observations classified only to the order level. The search for each family and the order Araneae were performed separately due to the limit that iNaturalist places on the amount of observation metadata that can be downloaded per query. The limit is 200,000 observations. Even by separating taxon into families, iNaturalist queries cannot return results with over 200,000 observations. For queries that exceed the 200,000-observation limit, modifications were made to one of the following criteria: Verifiable, Threatened, Introduced, Native, and Popular. Criteria in these selections can be selected as either “Any,” “Yes,” or “No.” By selecting “Yes” in one criterion, downloading the results, and then running the same query again but changing the criterion selection to “No,” the limit was no longer exceeded. Other selections are detailed in Figure 2. I obtained GBIF occurrence coordinates and image URLs using the Search Occurrences function for the order Araneae (“GBIF.org”).

**Figure 2.**
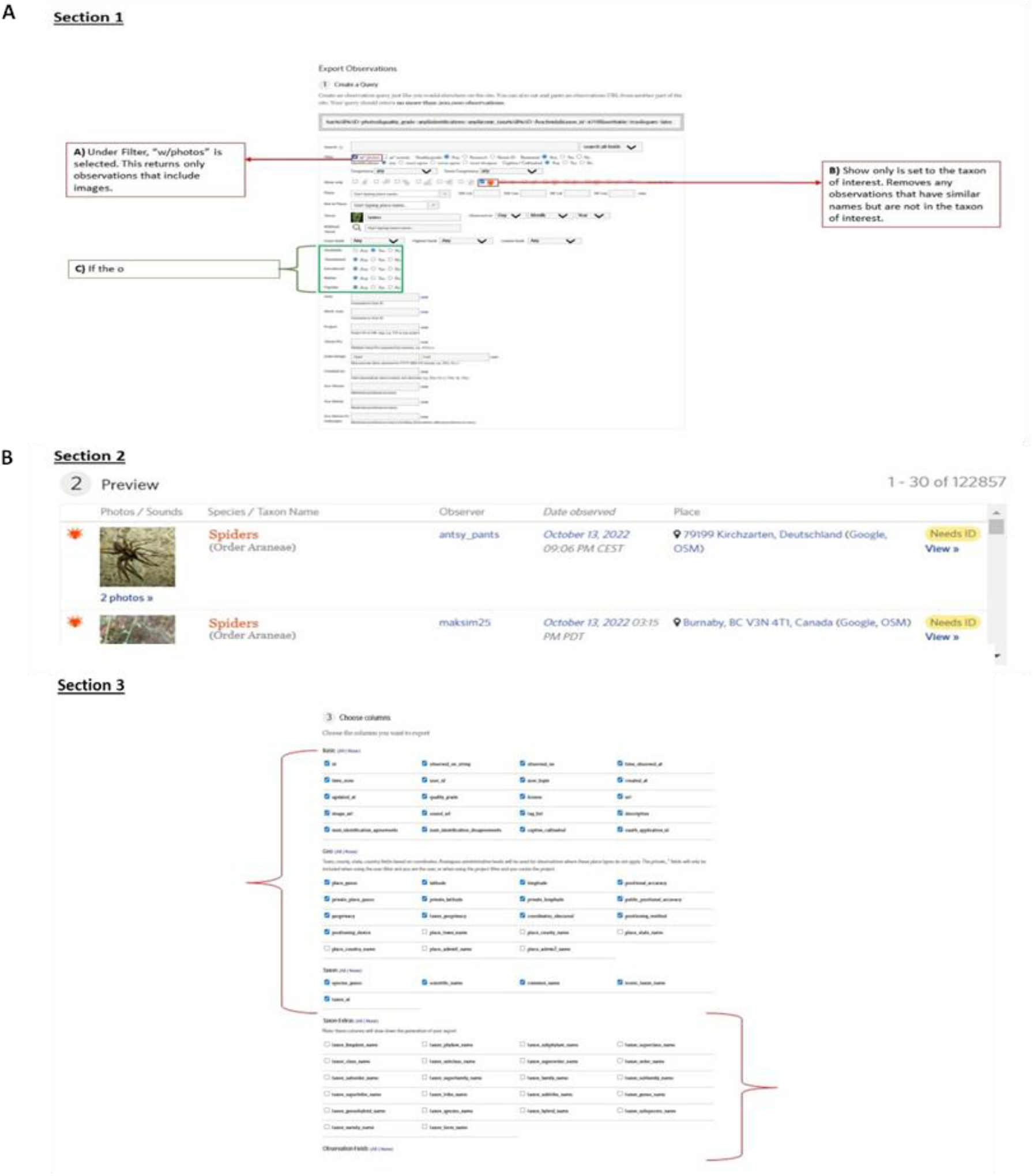
Option selections for downloading query results using the downloading function in iNaturalist. Part A) is the selection for Section 1. B) shows the selections for Section 3. No changes are made to the default settings in Section 2.

The pipeline uses an input file in a .*csv* format (a sample input file can be downloaded at https://doi.org/10.5281/zenodo.7352707 in the Sample Data file). The input file for iNaturalist observations and GBIF occurrences must include columns for observation ID, time of observation, date of observation, time zone, latitude, longitude, the images’ current grade label, the image URLs, taxonomic guess provided by the machine learning algorithm, and taxonomic labels at the family, genus, and species level. The input file is read into Rstudio for analysis. The resulting data frame for iNaturalist observations is queried for duplicate observations. Observations are considered duplicate entries if two observations have the same time of observation and location (Hochmairid et al., 2020).

iNaturalist does not currently provide a method that allows users to bulk download images. Instead, the image URLs are extracted from the input file into a separate URL list. Images are downloaded using the list of URLs with an original bash script included in the pipeline (RStudio, 2020).

### Taxonomic consistency

To evaluate the consistency of the taxonomic data from iNaturalist, a taxonomic reference file is needed. Several databases exist that can provide necessary taxonomic information on specific groups of animals, such as The Global Lepidoptera Names Index (Beccaloni et al., 2003), and The Mammal Diversity Database (Zenodo, 2022), and The Reptile Database (Uetz & Hallermann, 2021). This case study used an updated species list from the World Spider Catalogue (WSC) downloaded on July 30th, 2021, as a taxonomic reference. WSC collects arachnology literature and publications and maintains updated taxonomic lists (Natural History Museum Bern, 2019).

The input file for taxonomic reference should include columns for the highest classification being used in the pipeline, with consequent columns for all lower classifications (Supplemental 2). Taxonomic identification for the highest order of classification and all subsequent lower classifications are compared to the taxonomic reference file, respectively.

The percentage of observations that include taxonomic labels at each level of classification and the consistency of the assigned taxonomic names for each observation is then analyzed. If the identification given for an observation matches the recognized taxonomic names at the appropriate level and was consistent with accepted higher-order names as they appear on the taxonomic reference file, then it is recorded as correct. Correct labels are given one point per name. If the name on the observation is either entered incorrectly, entered at the incorrect taxonomic level (such as the genus taxonomy being recorded as the species), or does not align with the accepted higher order labels (i.e., the genus does not belong to the family assigned to the observation) it is recorded as incorrect (Figure 3). Incorrect or missing labels are given a score of zero. The awarded points are then divided by the number of taxonomic levels, to produce an overall score (Figure 3).

**Figure 3.**
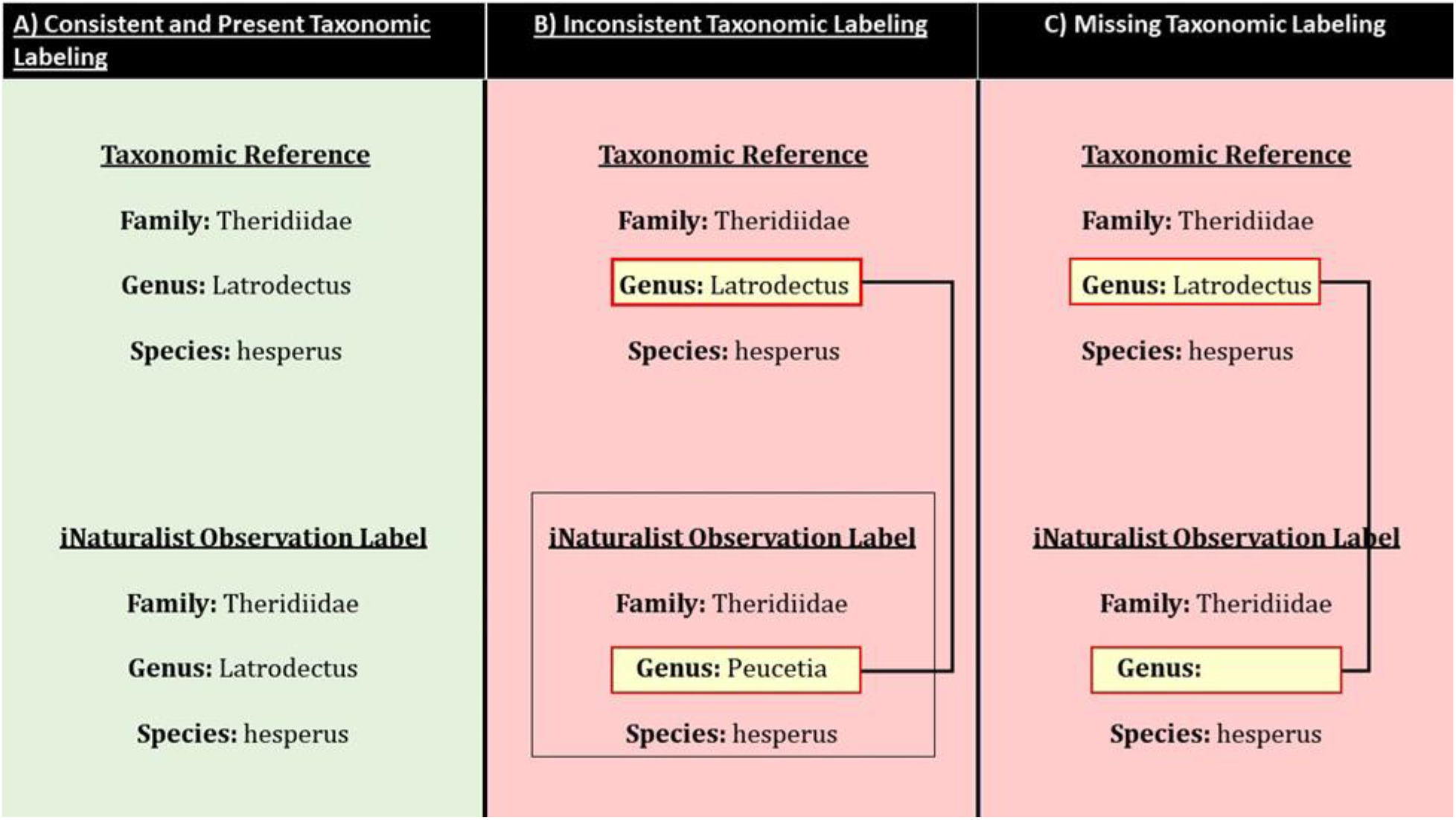
Taxonomic consistency workflow. Each observation could earn a maximum of three-points. Points were award as follows: iNaturalist observations were checked for family-level identification. Each level of taxonomic label (i.e. Family, Genus, and Species) was scored based on three criteria: 1) the label was present, 2) the label was an accepted name for that level of classification, 3) it agreed with all higher and lower taxonomic classifications. If these conditions were met, the observation was awarded an additional point. Lastly, observations were checked for species-level identification. To calculate the percent-correct the total number of observations positive for a level of identification was multiplied by three and then divided into the sum of all the scores for observations with each level of taxonomic identification.

### Geo-tagging

To assess geo-tagging data of observations, coordinates from iNaturalist and GBIF are filtered through the *Clean Coordinates* R-package, which removes coordinates with common errors found in biological and paleontological datasets (Zizka et al., 2019) These errors include locations at museum and zoo facilities, zero set coordinates, and coordinates that fall outside of any coordinate values, such as a latitude larger than 90°(Zizka et al., 2019).

Lists of the genera and species for observations from GBIF and iNaturalist are compiled and cross-referenced so only families, genera, and species that appear on both remain. A subset of coordinate data for both GBIF and iNaturalist are created, first by genera and then by species. Comparisons are made at the family, genus, and species levels. A Raster file is generated using GBIF occurrences in the R-program Raster and the raster polygon is divided into a grid GBIF and iNaturalist coordinates are converted to spatial points and placed on the grid. A confusion matrix is produced using the *confmat* function in the RStudio package GMDH2 (Dag et al., 2021). A confusion matrix is a type of contingency table that can be used as a performance test for classifications (Fawcett, 2006). on the results of each cell using the following parameters from Austen et al. (2018) (Figure 4):

**Figure 4.**
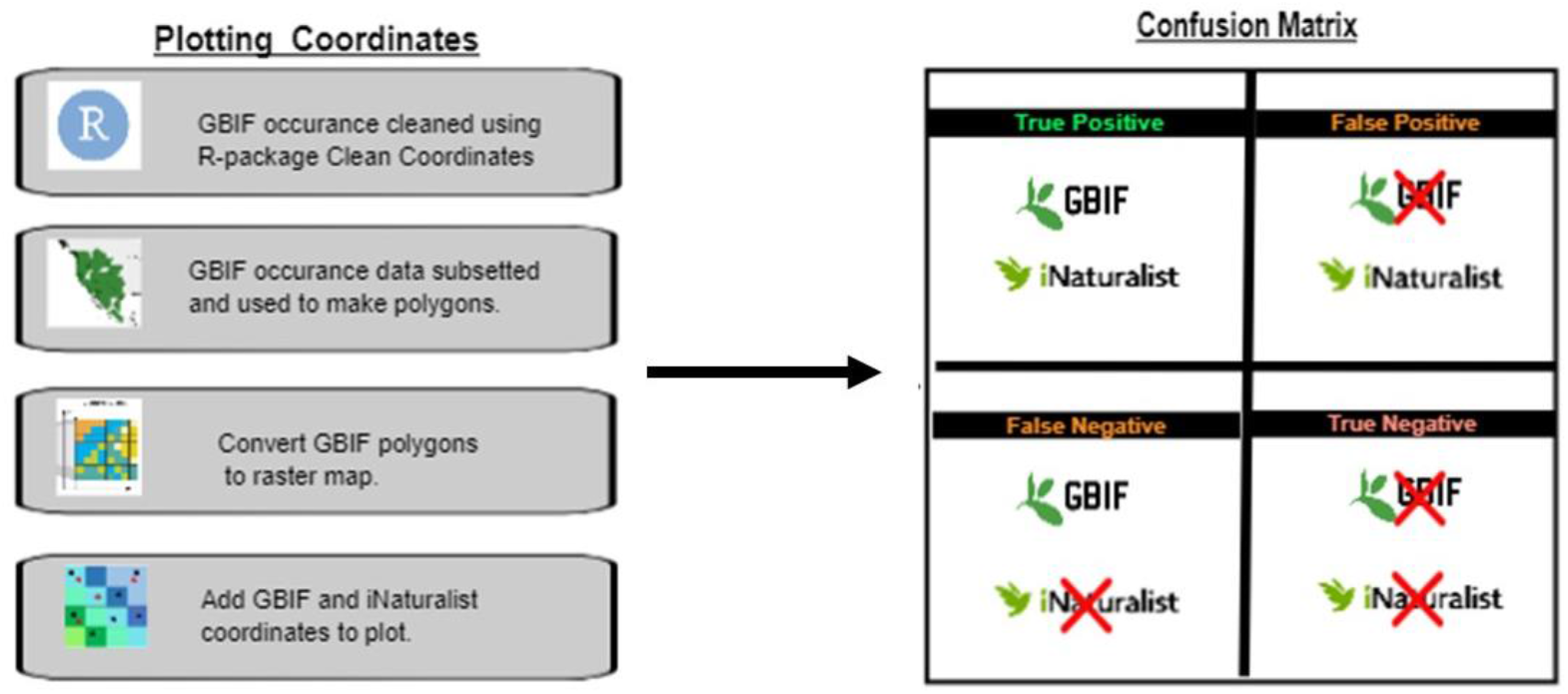
Geo-tagging Workflow. Plotting Coordinates: GBIF occurrence records was cleaned using the R-package, Clean Coordinates. Comparisons of occurrence records (including latitudes and longitudes) for iNaturalist observations were made using the family-level of identification, the genus-level of identification, and the species-level of identification. For each taxon rank, occurrence data was obtained from the Global Biodiversity Information Facility (GBIF) for the order Araneae and subsetted in Rstudio by taxon rank. The subsetted occurrence records were used to create polygon maps, that were then converted to raster maps. GBIF and iNaturalist occurrence records were added to the cells of the raster map. A confusion matrix was used to calculate the accuracy and precision of the iNaturalist occurrence records.

- True Positive: GBIF observation is present and iNaturalist
- True Negative: No observations present
- False Positive: iNaturalist observation present but not GBIF
- False Negative: GBIF observation present but not iNaturalist

Accuracy and precision scores from the confusion matrix are reported for each taxonomic level.

### Image quality

Image quality is assessed using the python version of the MatLab program BRISQUE, pybrisque (Gumbira, 2018). The BRISQUE algorithm scores the distortion in an image without the use of a reference image, to create a final quality score based on a comparison between the image and a default set of natural images (which are images captured directly from a camera without any post-processing) with similar distortions (Mittal et al., 2012). Images are evaluated and assigned a score that usually falls between 0 (higher quality) to 100 (lower quality) (Figure 5). Not all observations from iNaturalist included images with copyright permissions that allow them to be downloaded and some observations included multiple images, each with an individual score. Images are grouped by the highest shared taxonomic order and an average score is reported. This process is repeated for images with identification at lower taxonomic levels and a mean average score is calculated.

**Figure 5.**
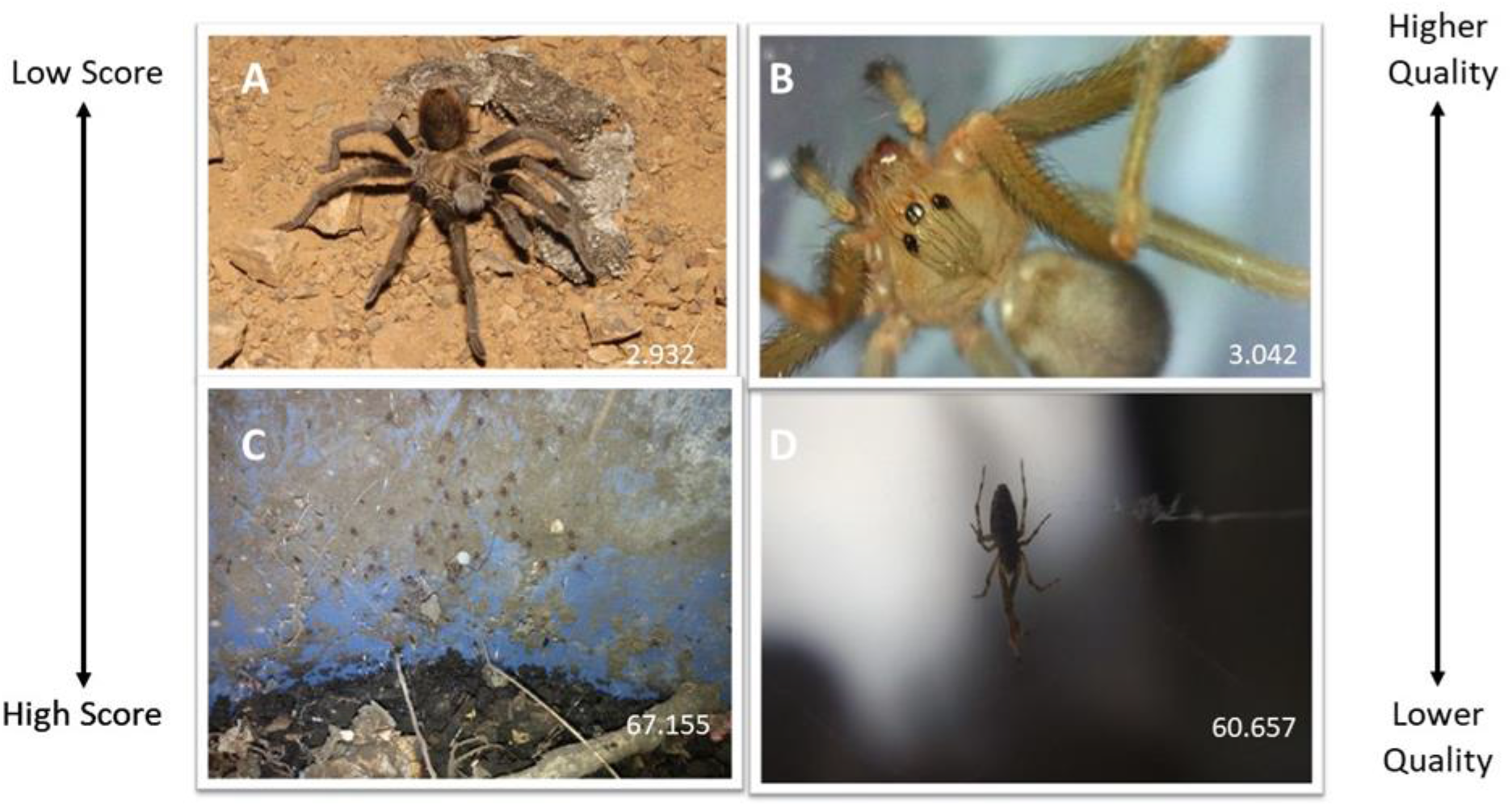
BRISQUE Image Quality. BRISQUE evaluates images without the need for a reference photo. The higher the score the lower the quality of the image. A comparison between images with a low BRISQUE score (A and B) and high BRISQUE scores (C and D). A was given a BRISQUE score of 2.932 and B was given a score of 3.042, while C was given a BRISQUE score of 67.155 and D was given a BRISQUE score of 60.657. A side-by-side comparison of these images demonstrates how visual assessment of image quality can be inaccurate and subject to individual preference and vision. Providing a BRISQUE score helps to quantify the quality of an image used. [A: Dunbar, 2014), B: (Miseroy, 2017), C: (Cuarenta, 2017), D: (veravilla, 2017).]

In addition to an overall average score, the image scores for the Araneae dataset are given at the family, genus, and species levels. To determine if a relationship existed between the level of taxonomic identification and the BRISQUE image score, I examined the normality of the data using Q-Q plots, residual plots, and a Shapiro-Wilks test. As the data were non-normally distributed, I used a Kruskal-Wallace test to compare the scores from the GBIF images, images with at most a family identification, genus identification, and species identification. All means are presented with a ±1 standard error of the mean.

## Results

### Data acquisition

For the order Araneae, I found 158,129 downloadable observations on iNaturalist. I found 78,310 unique observations for the order Araneae on iNaturalist. Of the 129 families in the order Araneae recognized by the WSC, 122 had observations (Natural History Museum Bern, 2019). I found no observations for Archoleptonetidae (though, as of August 2021 one observation has been added), Barychelidae, Huttoniidae, Mecicobothriidae, Myrmecicultoridae, Synaphridae, and Tetrablemmidae. Users of iNaturalist identified 49.91% of observations to at least the family level and so could be found searching by a specific family. Agelenidae (“Grass spiders”) had the most observations, with 41,716. A mean number of observations per family was 259.56 ±19.86. Archoleptonetidae and Penestomidae both had only one observation.

### Taxonomic consistency

I found 156,842 of the downloadable observations with a family-level identification, of which 79.15% were consistent with the taxonomic families listed on WSC. 99.86% of the 58,241 observations identified to the genus level and 99.74% of the 27,500 observations identified to the species level were consistent with established taxonomic names.

Observations may have more than one image associated with them, each with separate metadata. The metadata associated with each image is referred to as that image’s “record.” I removed 2,170 of 425,950 records after cleaning the coordinates. iNaturalist observations with a family-level identification were 49.95% accurate and 99.90% precise. In terms of the geo-tagged coordinates, “accuracy” refers to how often the coordinates from iNaturalist are present or absent in agreement with the coordinates from GBIF and are calculated using the following equation:

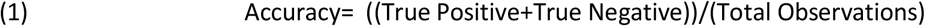

‘Precision” refers to the rate at which iNaturalist and GBIF coordinates are both present in cells on the grid, to the total number of grid scales in which iNaturalist coordinates are present:

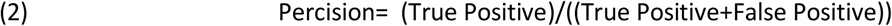

I found comparable results at the genus and species levels, with the genus level resulting in an accuracy of 49.97% and precision of 99.85%, and the species level resulting in an accuracy of 63.10% and precision of 99.87%.

### Image quality

pyBRISQUE evaluated 118,834 images. The median image quality score was 29.00 (n = 115,093, range = 70.32637). Images of Orsolobidae received the highest average BRISQUE score (n = 7, median= 36.95, range= 1.87), with a higher score indicating lower quality images. The family with the highest score and > 10 observations was Phyxelididae (n = 22, median = 36.94). Atracidae had the lowest (n = 176, median = 21.44, range = 26.92) and was also the family with the lowest score and > 10 observations.

## Discussion

Here, I present a method for assessing the metadata and image quality for data acquired from iNaturalist, a crowd-sourced biodiversity database. Using the order Araneae, both the metadata and images associated with observations in iNaturalist demonstrate that this platform provides a diverse dataset for spiders (Order Araneae). This data assessment method was able to determine that, with few exceptions, nearly every family in the order had some level of representation on iNaturalist. Automizing a check of taxonomic labels against accepted taxonomic names permits the quick removal of occurrences with inconsistent labels. Additionally, it provides researchers that use these data to report a quantifiable assessment of the taxonomic consistency in a dataset.

Similarly, comparing geo-tagging data from iNaturalist observations to GBIF occurrences allows for a quantifiable evaluation of that data via the confusion matrix results. Another benefit to using this pipeline to compare geo-tagging data is the insight into the accuracy of the observations’ taxonomic labels it can provide. Observations that fall outside the raster generated by the occurrence data from GBIF can be seen as outside the area where that organism has previously been shown to live. This could also aid in determining if the iNaturalist taxonomic labels are accurate. Conversely, this method of evaluating geo-tagging could help identify if a species was observed within a new range. While an occurrence could fall outside of the GBIF raster for reasons unrelated to range expansion, once the raster polygon is created using the GBIF occurrences, iNaturalist observations outside of the raster are easily separated from the dataset and further investigated.

One major concern with using publicly sourced biodiversity photos is that without knowing the quality of the images, reproducibility is difficult. iNaturalist categorizes images as either “research grade,” “needs ID,” or “NA.” iNaturalist’s current method for assigning an observation as “research grade” or not “research grade” requires that an observation reach a two-thirds consensus from users to confirm identification (Agrin et al., 2014). Images currently marked “needs ID” could receive the consensus on their taxonomic idea and be relabeled as “research grade.” I found relatively consistent image quality across families and between research grade and non-research grade observations (“needs ID” and “NA”). While this system of categorization does provide some information on the quality of an image, presenting a mean score for image sets on databases may help clarify the results researchers obtain using iNaturalist (or another database) images, especially in the cases of machine learning programs. The method presented for image assessment could also be used to set a threshold score, allowing only images that meet a specific quality standard to be included in a set of images.

Additionally, this study highlights the importance of using quantifiable image quality assessment tools like BRISQUE, especially for methods that use images for morphological evaluation of computer learning software for identification. While not conducted here, it would be beneficial to perform a BRISQUE assessment on any image used for research and to report the score or average score of images, especially when sourced from public collections. Image quality could impact evaluations of coloring and morphological structures. From a visual assessment of an image, the quality of that image may not be obvious. Additionally, visual assessment of an image may be subject to personal bias or visual acuity (Wang et al., 2004). Providing a quantitative score, such as a BRISQUE does, permits a uniform understanding of the images being used. In the future, it would be beneficial to compare the BRISQUE scores of images obtained from iNaturalist to those on image databases that only host published images, such as GBIF.

One drawback to using crowd-sourced observations, however, is how usage patterns could affect the metadata and number of high-quality images available for a specific group of animals. For example, differences in the number of observations per family of Araneae might result from geographic areas with a higher or lower user number of iNaturalist users, variation in numbers of people with access to camera phones or reflect actual relative abundances.

Increasing the number of observations at the family level or lower is the best way to increase the number of images and volume of metadata available to researchers from iNaturalist, since a lack of identification, lack of consensus for that identification, or no lower order identification represents the most common deficits for most observations. To help increase the diversity and quality of the observations, researchers should employ methods to contribute to iNaturalist, such as educational initiatives that engage students with the platform and encourage them to contribute images or by making a habit of offering taxonomic identifications to images that require taxonomic identification.

For researchers to become more engaged with iNaturalist, however, some changes may be necessary. For example, limits on how many observations could be downloaded at one time make broad studies, such as the one using Araneae, more time-consuming than necessary. iNaturalist has recently launched an Amazon Web Services (AWS) platform, but this platform does not include the image files from the database, which limits the type of analyses researchers can conduct. In addition to a more economical method to bulk download images, incorporating an image assessment algorithm like BRISQUE into the metadata would increase the utility of iNaturalist to researchers. Providing a BRISQUE score would allow users to filter downloads for a specific image quality without having to run the algorithm on large image datasets. Finally, while not an issue for iNaturalist observations, problems exist in downloading images from GBIF in bulk and keeping the image file names associated with the taxon. Another database may experience similar problems.

Several databases are available for researchers to access occurrence data and images(Shirey et al., 2019; Heberling et al., 2020; Moudrý & Devillers, 2020; Cull, 2021). iNaturalist provides a user-friendly, crowd-sourced database of images with the potential for broad research applications (Hochmairid et al., 2020; Nugent, 2020; Mesaglio & Callaghan, 2021). It should be noted that any research that uses images or metadata from iNaturalist should include methods that screen the data for duplicates, incorrectly geo-tagged observations, and image quality.

The usefulness of this pipeline may be limited for systems with few observations on iNaturalist or GBIF (who must have at least three occurrences to generate a raster polygon). Systems that lack a taxonomic database could also present a challenge since this would require the user to generate the taxonomic reference file from scratch. An alternative to creating a *de novo* taxonomic reference is sourcing taxonomic names from an established database such as the Integrated Taxonomic Information System (“Integrated Taxonomic Information System”). GBIF also provides a taxonomic database. More specialized taxonomy databases could also serve as a source of taxonomic comparison. The Mammal Diversity Database is maintained by the American Society of Mammologists and provides an updated list of known mammal taxonomic names and species. Similar initiatives exist for amphibians, ants (AntWeb), birds (Avibase, IOC World Bird List), fungi, and reptiles. General and specialized databases could be cross-referenced or used independently to generate the reference needed to assess taxonomic consistency.

Taken together, this study demonstrates that iNaturalist can provide large metadata and image datasets for research. This pipeline can be used to assess the taxonomic consistency, relationship to known distributions, and image quality of large datasets of crowd-sourced data. This is a reliable method to quickly analyze the data quality for specific taxa. With appropriate quality controls in place, the wealth of knowledge supplied through crowd-sourced biodiversity databases can more reliably be used for scientific discovery.

## Supporting information

iNaturalist Metadata

Sample Taxon Reference

Sample GBIF Metadata

Sample GBIF Image URLs

## Acknowledgments

I would also like to acknowledge and thank the reviewers of this paper, Catherine Scott and Clive Hambler for their thoughtful comments and efforts in improving our manuscript. I would like to thank Ruth Hufbauer, Richard Reading, and Lorna McCallister. for providing comments on this manuscript. Preprint version 5 of this article has been peer-reviewed and recommended by Peer Community In Zoology (https://doi.org/10.24072/pci.zool.100017).

## Data, scripts, code, and supplementary information availability

Data are available online: https://doi.org/10.5281/zenodo.7352707

Scripts and code are available online: https://doi.org/10.5281/zenodo.7352707

